# Mechanisms of ligand recognition and activation of melanin-concentrating hormone receptors

**DOI:** 10.1101/2023.11.03.565472

**Authors:** Qian He, Qingning Yuan, Hong Shan, Canrong Wu, Yimin Gu, Kai Wu, Wen Hu, Yumu Zhang, H. Eric Xu, Li-Hua Zhao

## Abstract

Melanin-concentrating hormone (MCH) is a cyclic neuropeptide that regulates food intake, energy balance, and other physiological functions by stimulating MCHR1 and MCHR2 receptors, both of which are class A G protein-coupled receptors. MCHR1 can couple with multiple G-proteins, including G_i/o_, G_q/11_, and G_s_, while MCHR2 only couple to G_q/11_. Here we present cryo-electron microscopy structures of MCH-activated MCHR1 with G_i_ and MCH-activated MCHR2 with G_q_ complexes, at the global resolutions of 3.01 Å and 2.40 Å, respectively. These structures reveal that MCH adopts a consistent cysteine-mediated hairpin loop configuration in both receptors. A central arginine from the LGRVY core motif between the two cysteines of MCH penetrates deeply into the transmembrane pocket, triggering receptor activation. Integrated with mutational and functional insights, our findings elucidate the molecular underpinnings of ligand recognition and MCH receptor activation, offering a structured foundation for targeted drug design.

---

Melanin-concentrating hormone (MCH) is a cyclic neuropeptide of 19 amino acids, predominantly synthesized by neurons in the hypothalamus and the zona incerta of the brain ^1–5^. Originally identified in fish due to its role in melanin aggregation within skin melanophores, MCH in mammals orchestrates a myriad of physiological functions ^1^. These range from energy homeostasis, appetite regulation, and sleep-wake cycles to mood modulation, stress responses, and reproductive functions^6–14^. Disruptions in MCH signaling pathways have been associated with obesity, psychiatric conditions, sleep disorders^15–19^. Mice lacking MCH system display a lean phenotype, diminished appetite, increased mobile activity, and metabolic shift^20–23^. Consequently, the MCH system is increasingly recognized as a promising therapeutic target for conditions like obesity, depression, and sleep disorders.

MCH exerts its effects through two specific G protein-coupled receptors: MCHR1 and MCHR2, both predominantly found in the central nervous system ^24^. The binding of MCH to these receptors triggers conformational changes, initiating intracellular signaling cascades via heterotrimeric G proteins. Specifically, MCHR1 couples with G_i/o_, G_q/11_, and G_s_ proteins^24–27^, whereas MCHR2 is exclusively coupled to G ^24,28^ (Fig. 1a). While MCHR1 antagonists are primarily explored for obesity treatment, they also show promise in addressing anxiety and depression ^19,29–33^. However, a challenge arises as some antagonists interact with the hERG (human *ether-a-go-go-related gene*) channel, leading to cardiovascular complications^31^. To design drugs with enhanced specificity and minimal off-target effects, a comprehensive understanding of the structural mechanisms of MCH receptor activation is paramount. In this paper, we present cryo-EM structures of MCH-activated MCHR1-G_i_ and MCHR2-G_q_ complexes, with resolutions of 3.01 Å and 2.40 Å, respectively. These structures shed light on the molecular intricacies of ligand binding, G protein coupling, and receptor activation, paving the way for structure-based drug design targeting MCH receptors.

**Fig. 1.**
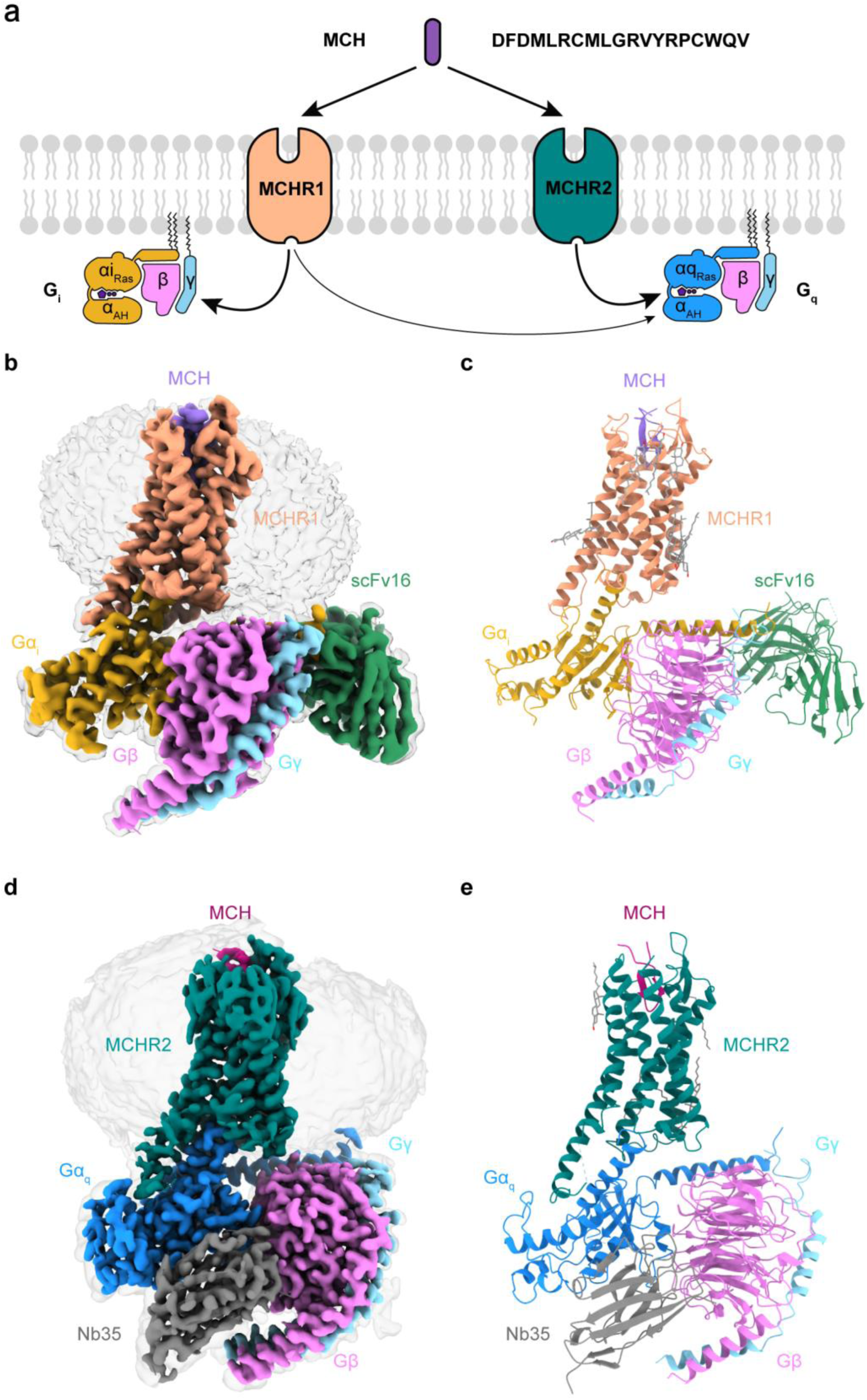
Structural representations of the MCH-MCHR1-G_i_ and MCH-MCHR2- G_q_ complexes. **a**, Depiction of the MCH sequence and a schematic illustrating the G-protein coupling of MCHR1 and MCHR2. **b**,**c**, Cryo-EM density map (**b**) and the corresponding molecular model (**c**) of the MCH-MCHR1-G_i_ complex. **d**,**e**, Cryo-EM density map (**d**) and the corresponding molecular model (**e**) of the MCH-MCHR2-G_q_ complex.

## Structure determination and overall structures

Our group has screened expression of many GPCRs, both human MCHR1 and MCHR2 present extremely challenges in their protein expression. To enhance cell-surface expression, a truncation of 69 amino acids from the N-terminus of MCHR1 was carried out, as previously reported^5^. Additionally, a BRIL was fused to the N-terminus of both MCHR1 and MCHR2^34^. To further stabilize the complex, we employed the NanoBiT tethering strategy, fusing the large BiT (LgBiT) to the C-terminus of MCHR1 (residues 70-405) and MCHR2 (residues 1-331), which would form a complex with small BiT (HiBiT) attached to the C-terminus of Gβ as detailed below^35–37^. A dual maltose-binding protein (MBP) tag was appended after the LgBiT to augment receptor expression and streamline protein purification ^38^.

For the assembly of the MCH-MCHR1-G_i_ complex, MCHR1, Gα_i_, Gβγ subunits, Ric8B, and scFv16 were co-expressed in Hi5 insect cells and subsequently incubated with MCH. Conversely, the MCH-MCHR2-G_q_ complex was formed by co-expressing MCHR2, Gα_q_, Gβγ subunits, and Ric8A in Hi5 insect cells, followed by MCH incubation. The Gα_i_ in the MCHR1 complex harbors dominant-negative mutations (S47N, G203A, A326S, E245A) ^39^, which diminish nucleotide-binding affinity and stabilize the Gαβγ heterotrimer complex. The Gα_q_ in the MCHR2 complex is a mini- Gα_q_ variant, integrating the C-terminus of Gα_q_ with a mini-Gα_s_ framework. This configuration permits the addition of nanobody (Nb35) to bolster the stability of the receptor-G protein complex. The Gβ subunit in both MCHR1 and MCHR2 complexes contains a C-terminal HiBiT, which interacts with the LgBiT at the C-terminus of receptors, effectively anchoring the G protein complex to the receptor.

The structures of the MCH-MCHR1-G_i_ and MCH-MCHR2-G_q_ complexes were resolved by cryo-EM to 3.01 Å and 2.40 Å, respectively, as depicted in Fig. 1b-e and Extended Data Fig. 1, 2. The clarity of the cryo-EM density maps allowed for the accurate modeling of most side chains, encompassing MCH, the receptors, the Gαβγ heterotrimer, and both scFv16 and Nb35(Fig. 1c,e and Extended Data Fig. 3). In terms of receptor composition in the final structure models, MCHR1 spans residues 107 to 393, with an absent ICL1 segment (residues 139-142). MCHR2, on the other hand, covers residues 31 to 317, but lacks ICL3 fragments from positions 242 to 244. Regarding the ligand, the residues 2-18 of MCH in the MCH-MCHR1-G_i_ complex is well-defined. However, in the MCH-MCHR2-G_q_ complex, the N-terminal 4 amino acids of MCH are absent from the EM map. In addition, 8 cholesterol molecules were found to surround the MCHR1 transmembrane domain (TMD) but only 3 cholesterol molecules were found in the MCHR2 structure (Fig. 1c, e).

The overall structures of MCHR1 and MCHR2 in both complexes are highly similar, adopting a canonical seven transmembrane helix domain (TMD), with root mean square deviation (RMSD) values of 1.02 Å for the receptors and 1.86 Å for the complete complexes. The ligand, MCH, also adopts a similar β-hairpin structure in both receptor complexes, with an RMSD value of 0.51 Å for the central 10 residues between the two disulfate-bond cysteines (C7 and C16). The binding site of MCH in both receptors is at the orthosteric pocket formed by transmembrane helices TM2, TM3, and TM5-7, as well as three extracellular loops (ECL1-3). The overall binding mode of MCH to MCHR1 and MCHR2 is similar to the binding mode of other cyclic peptides, including somatostatin-14 (SST14) to somatostatin receptor 2 (SSTR2) and arginine vasopressin (AVP) to vasopressin type 2 receptor (V2R) (Fig. 7 and Extended Data Fig. 4).

## MCH recognition by MCHR1

MCH in the MCHR1 complex adopts a γ-shape configuration, with the central 10 residues forming a cyclic loop with a disulfate bond by cysteine residues at positions 7 and 16. The central loop residues are buried in the TMD pocket, with the central arginine from the conserved LGRVY core motif (residues 9-13) inserted deeply into the bottom of the pocket. The N-terminal portion of MCH (residues 2-6) runs toward TM2 and ECL1, where the C-terminal portion of MCH (residues 17-18) runs towards TM5 and packs along with ECL2 (Fig. 2a).

**Fig. 2.**
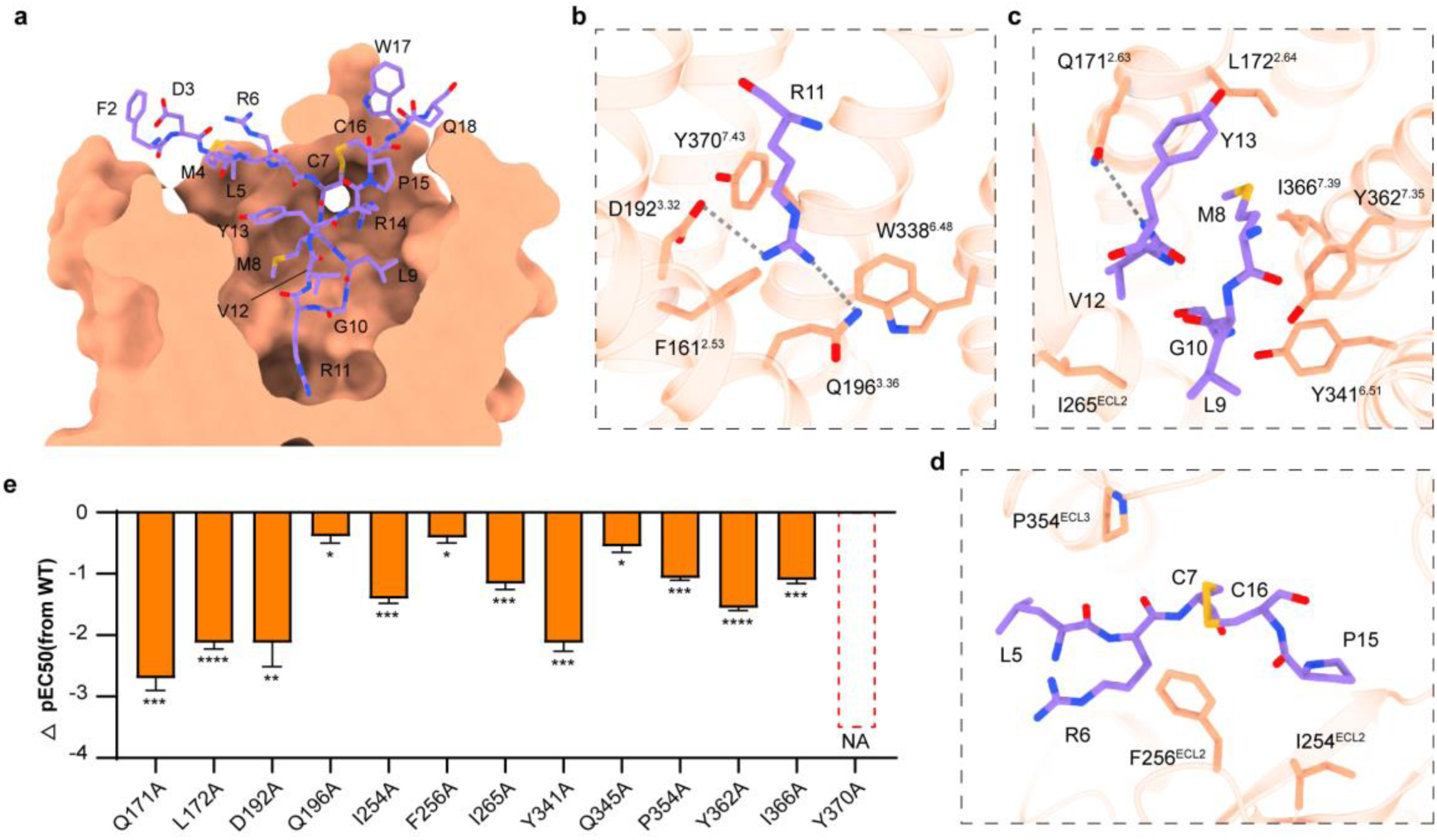
Molecular basis of MCH recognition by MCHR1. **a**, Sequence representation of MCH alongside a side-view of the MCHR1 binding pocket. **b**-**d**, Detailed molecular interactions between MCH (depicted in medium purple) and MCHR1 (shown in light salmon). Hydrogen bonds are presented with gray dashed lines. **e**, Functional implications of mutations within the MCHR1 binding pocket, represented as ΔpEC50 values. ΔpEC50 is derived by subtracting the pEC50 of mutations from the pEC50 of the wild-type (WT) MCHR1. Data are presented as mean ± S.E.M. from three independent experiments performed in triplicate and analyzed by two-sided unpaired t-test. NA, no activity; *p < 0.05; **p < 0.01; ***p<0.001; ****P < 0.0001.

The binding mode of MCH in MCHR1 is stabilized by extensive interactions of MCH with specific residues in the receptor binding pocket, as summarized in Table 1. The most prominent interaction is mediated by R11 of MCH, which side chain is docked near the bottom of the TMD pocket and forms an electrostatic interaction with D192^3.32^, a hydrogen bond with Q196^3.36^ (Fig. 2b). Surrounding the central arginine are several hydrophobic residues (9-LGRVY-13), which form extensive hydrophobic interactions with residues F161^2.53^, W338^6.48^, Y370^7.43^, L172^2.64^, I265^ECL2^, Y341^6.51^, Y362^7.35^ in the TMD pocket (Fig. 2b,c). Besides, the amino group of Y13 main chain forms a hydrogen bond with Q171^2.63^ (Fig. 2c). Other hydrophobic residues within the cyclic loop (M8 and P15) also make hydrophobic interactions with receptor residues (Y362^7.35^, I366^7.39^ and I254^ECL2^) in the upper part of the TMD pocket (Fig. 2c,d). In addition, the N-terminal hydrophobic residues L5 contacts residues from ECL2 and 3, including P256^ECL2^, P354^ECL3^ (Fig. 2d). Alanine mutation at key pocket residues resulted in varied degrees of reduced receptor activation, with the Y370^7.43^ mutation resulting in complete loss of activity, highlighting the importance of these interactions (Fig. 2e and Extended Data Fig. 5).

## MCH recognition by MCHR2

MCH recognition mode by MCHR2 closely resembles that in MCHR1, with the MCH displaying a similar binding pose. The central arginine from the conserved LGRVY core motif also is deeply buried into the bottom of the orthostatic pocket of MCHR2 (Fig. 3a). Overall, MCH is make very similar interactions with MCHR2 and MCHR1, as summarized Table 1-3 and Fig. 3b-f. However, MCH makes more extensive and tighter interactions with the residues from MCHR2, especially with ECL2 of MCHR2 (Fig. 3c-f).

**Fig. 3.**
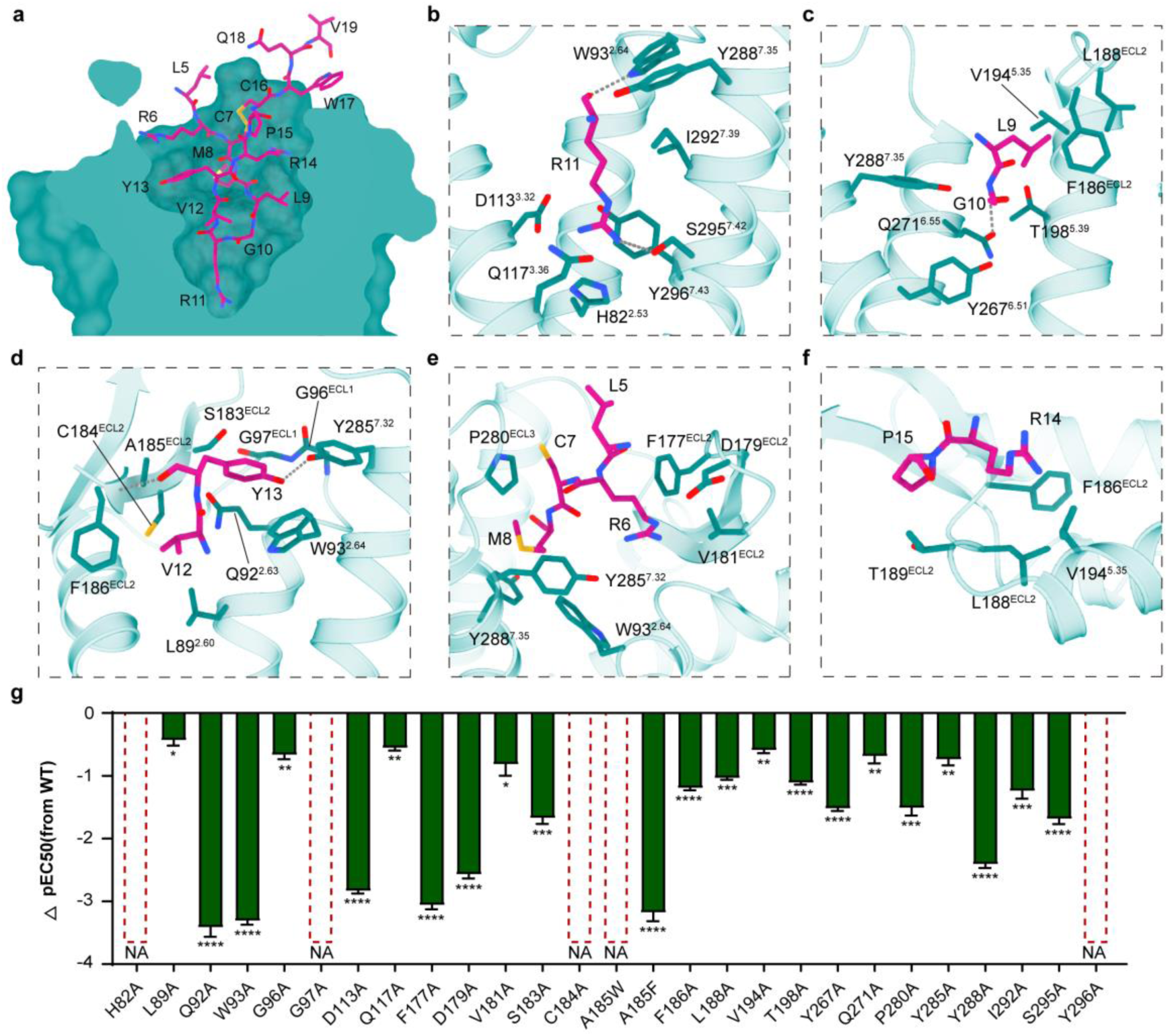
Molecular basis of MCH recognition by MCHR2. **a**, Sequence representation of MCH and a side-view of the MCHR2 binding pocket. **b**-**f**, Detailed molecular interactions between MCH (depicted in medium violet red) and MCHR2 (shown in teal). Hydrogen bonds are highlighted with gray dashed lines. **g**, Functional data derived from alanine mutations of residues within the MCHR2 binding pocket, represented as ΔpEC50 values. ΔpEC50 is derived by subtracting the pEC50 of mutations from the pEC50 of the wild-type (WT) MCHR2. Data are presented as mean ± S.E.M. from three independent experiments performed in triplicate and analyzed by two-sided unpaired t-test. NA, no activity; *p < 0.05; **p < 0.01; ***p<0.001; ****P < 0.0001.

In addition to various similar interactions, MCH and MCHR2 engage in additional polar interactions. The central arginine of MCH can form hydrogen bonds with W93^2.64^, S295^7.42^ of MCHR2 (Fig. 3b). The corresponding positions of MCHR1 are substituted by L172^2.64^ and G369^7.42^, which cannot form hydrogen bonds with R11. Among the other residues of core motif, the backbone amine group of G10 and carbonyl group of Y13 can form hydrogen bonds with Q271^6.55^ and the main chain amine group of F186^ECL2^, respectively, and both of the hydroxyl group of Y13 and Y285^7.32^ form a hydrogen bond (Fig. 3c,d). The sidechain of R6 form electrostatic interaction with D179^ECL2^ (Fig. 3e). In addition, ECL2 of MCHR2 forms a hydrophobic cap composed of F177^ECL2^, V181^ECL2^, S183^ECL2^, C184^ECL2^, A185^ECL2^, F186^ECL2^, L188^ECL2^, which makes intensive hydrophobic interactions with MCH residues L5, L9 and V12-P15 (Fig. 3c-f).

Consistently, mutations of the receptor pocket residues above to alanine significantly decrease or even completely abolished receptor activity, underscoring the essential role of these amino acids in the binding of MCH to MCHR2 (Fig. 3g and Extended Data Fig. 5). In addition, MCH is closely packed against ECL2 of MCHR2 (Fig. 3c-f). The A185^ECL2^F mutation results in a decrease in activity by more than 1000-fold, while the A185^ECL2^W mutation completely abolishes activity (Fig. 3g). This suggests that increased steric hindrance of ECL2 may hinder MCH from effectively accessing the ligand-binding pocket.

## Differences in MCH binding between MCHR1 and MCHR2

While MCHR1 and MCHR2 share highly similar overall structures and MCH binding modes, subtle differences in their binding pockets confer differences for MCH (Fig. 4a-c). Notably, residues deep within the binding pocket of MCHR2 make more extensive interactions with the critical R11 of MCH. R11 forms hydrogen bonds with S^7.42^ and W^2.64^ in MCHR2, enabling a unique interaction pattern (Fig. 4d). In contrast, in MCHR1, R11 shifts away from TM6 due to a potential steric clash with W^6.48^ and the absence of a hydrogen bond with G^7.42^ (corresponding to A^6.48^ and S^7.42^ in MCHR2) (Fig. 4d).

**Fig. 4.**
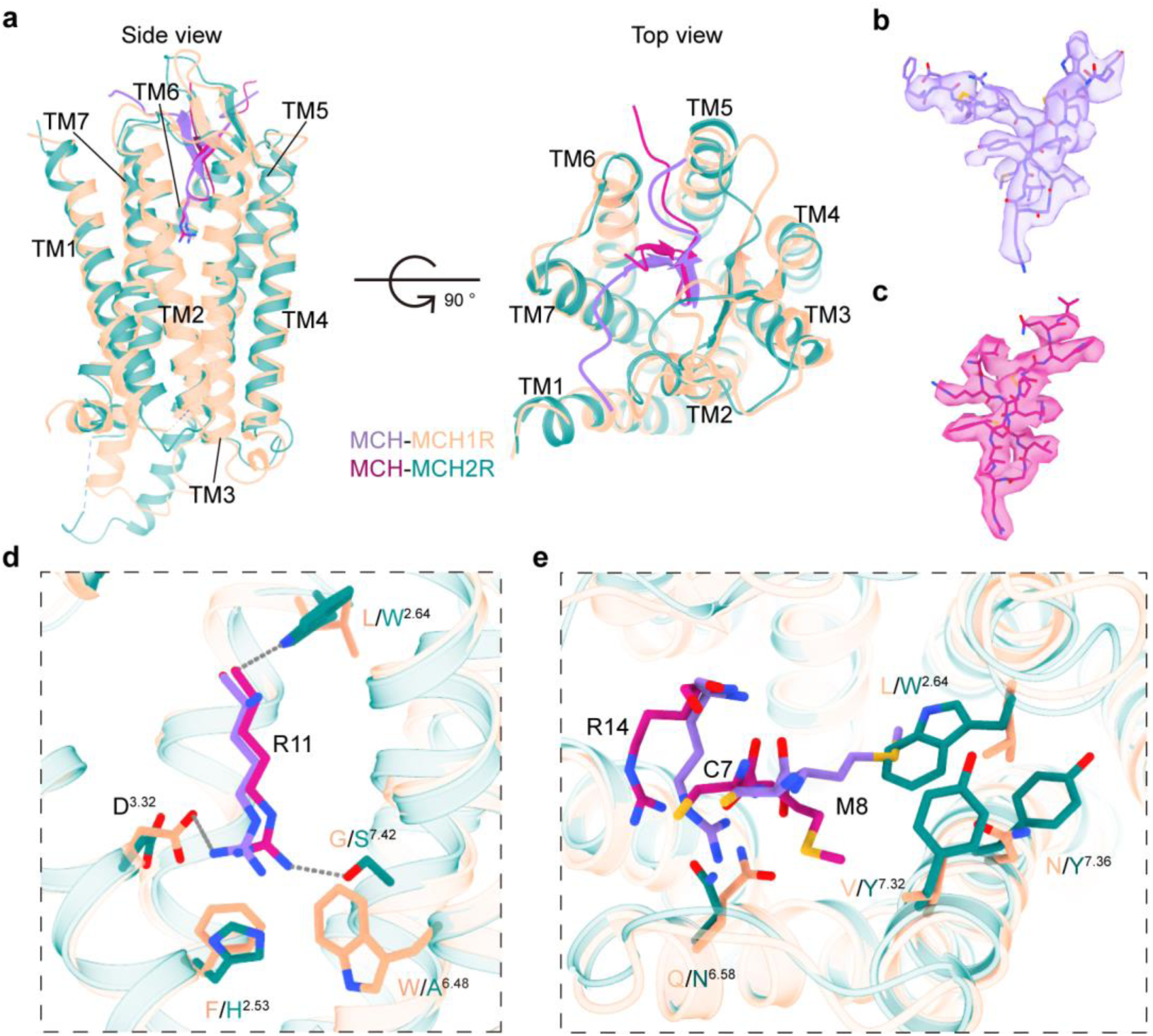
Conformational divergence of MCH recognition by MCHR1 and MCHR2. **a**, Structural superposition of MCH-bound MCHRs, presented from both side-view (the left) and top view (the right). **b**,**c**, Mesh representations of MCH within the binding pockets of MCHR1(**b**) and MCHR2(**c**). **d**,**e,** Highlighting the differential interactions between MCH and MCHRs, focusing on R11 (**d**) and the C7, M8, R14 residues of MCH (**e**).

Additionally, residues 2.64 and 7.32 confer other differences. MCHR1 has L^2.64^ and V^7.32^, while MCHR2 has W^2.64^ and Y^7.32^. The bulky tryptophan and tyrosine residues introduce steric hindrance that brings C7, M8, and L9 of MCH closer to TM4 and TM5 in MCHR2 (Fig. 4e). This enhances interactions between MCH and the extracellular loop 2 (ECL2) of MCHR2, which dips slightly towards the binding pocket. Furthermore, R14 of MCH clashes with N^6.58^ in MCHR1, causing it to shift downwards compared to Q^6.58^ in MCHR2 (Fig. 4e). In concert, these distinctions display unique differences in MCH binding to MCHR1 from MCHR2.

## Activation mechanism of MCHRs

To elucidate the activation mechanisms of MCHR1 and MCHR2, we compared their active structures to the inactive SSTR2 (PDB: 7UL5), which shares the highest homology with MCHRs among all GPCRs ^25,40^(Fig. 5a). Structural alignment showed that upon MCH binding and activation, the cytoplasmic ends of TM6 in MCHR1 and MCHR2 moved outward by approximately 9.7 Å and 11.0 Å respectively, as measured at the Cα of residue 6.30 (Fig. 5b). This outward movement of TM6 is a hallmark of GPCR activation and enables coupling to downstream G proteins. Additionally, TM7 shifted inward by 2.6 Å in MCHR1 and 1.5 Å in MCHR2 at the Cα of residue 7.53 (Fig. 5b). The combination of TM6 and TM7 movements opens a cleft on the intracellular side to accommodate G protein binding.

**Fig. 5.**
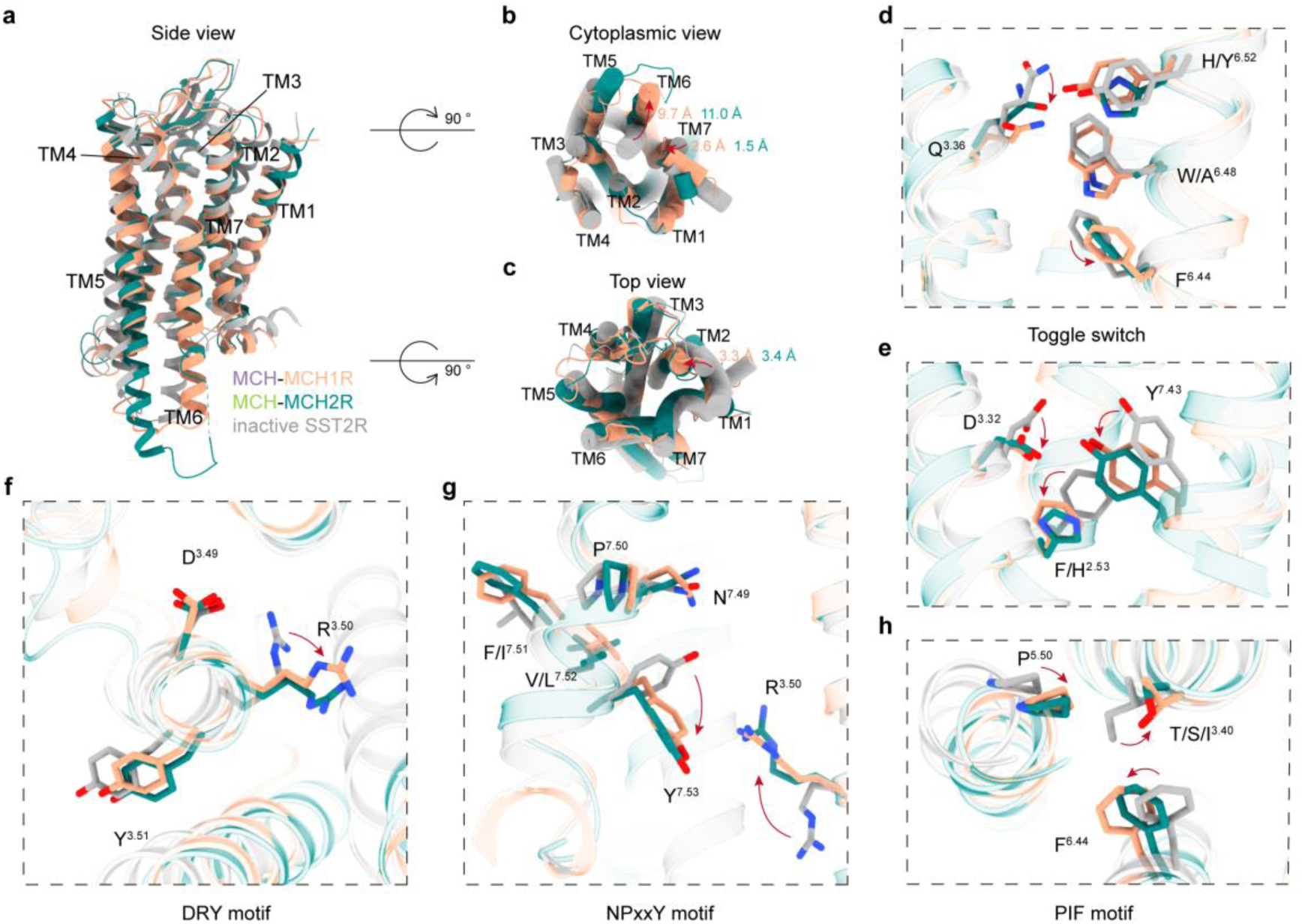
Conformational changes in MCHR1 and MCHR2 upon activation. **a**,**b**,**c**, Structural superposition between MCH-bound MCHRs and the inactive SSTR2 (PDB: 7UL5) viewed from the side view (**a**), cytoplasmic view (**b**), and top view (**c**). **d**, Structural alignment highlighting the toggle switch mechanism. **e**, Depiction of shifts in H/F^2.53^, D^3.32^, and Y^7.43^ at the base of the binding pocket. **f**-**h**, Illustration of the conserved motifs in MCH-activated MCHRs: (**f**) DRY motif; (**g**) NPxxY motif; (**h**) PIF motif.

The receptor activation is induced by MCH binding, which inserts into the orthosteric binding pocket to induce conformational changes that propagate through the helices (Fig. 5c). Notably, most class A GPCRs contain a highly conserved toggle switch W^6.48^ that shifts downwards upon activation. However, in MCHR1 this tryptophan shows minimal movement, and MCHR2 contains an alanine substitution at this position (Fig. 5d). Instead, in both receptors, MCH disrupts the polar interaction between Q^3.36^ and H/Y^6.52^, causing displacement of these residues (Fig. 5d).

Furthermore, R11 of MCH forms critical interactions with D^3.32^ and Y^7.43^ at the base of the pocket. This results in downward shifts of D^3.32^ and Y^7.43^, which in turn induce the upwards movement of R^3.50^ in the DRY motif and subsequent downward shift of Y^7.63^ in the NPxxY motif (Fig. 5e-g). The H/Y^6.52^ displacement also shifts F^6.44^ in the PIF motif (Fig. 5h). Collectively, these rearrangements within key structural motifs facilitate the activated state.

## G-protein coupling of MCHR1 and MCHR2

MCHR1 and MCHR2 exhibit distinct G-protein coupling profiles. Specifically, MCHR1 is capable of associating with a diverse set of G-proteins, namely Gα_i/o_, Gα_q/11_, and Gα_s_. Conversely, MCHR2 demonstrates specificity by predominantly coupling to Gα_q/11_. Experimental analyses further elucidated that MCHR1 manifests a heightened efficacy in its coupling to Gα_i_ as opposed to Gα_q_, as depicted in Extended Data Fig. 5c.

Structural alignments between Gα_i_ and Gα_q_ have unveiled notable conformational disparities (Fig. 6a). The α5 helix of both Gα_i_ and Gα_q_ exhibits a deflection of approximately 7°, and a discernible spatial separation of ∼10.3 Å is observed in their respective αN regions (Fig. 6b,c). Intriguingly, both MCHR1 and MCHR2 engage with Gα_i_ or Gα_q_ through transmembrane domains 2 to 7 (TM2-7) and intracellular loops 2 and 3 (ICL2 and ICL3) (Fig. 6d). An additional interaction via TM1 is uniquely observed in MCHR2 (Fig. 6d).

**Fig. 6.**
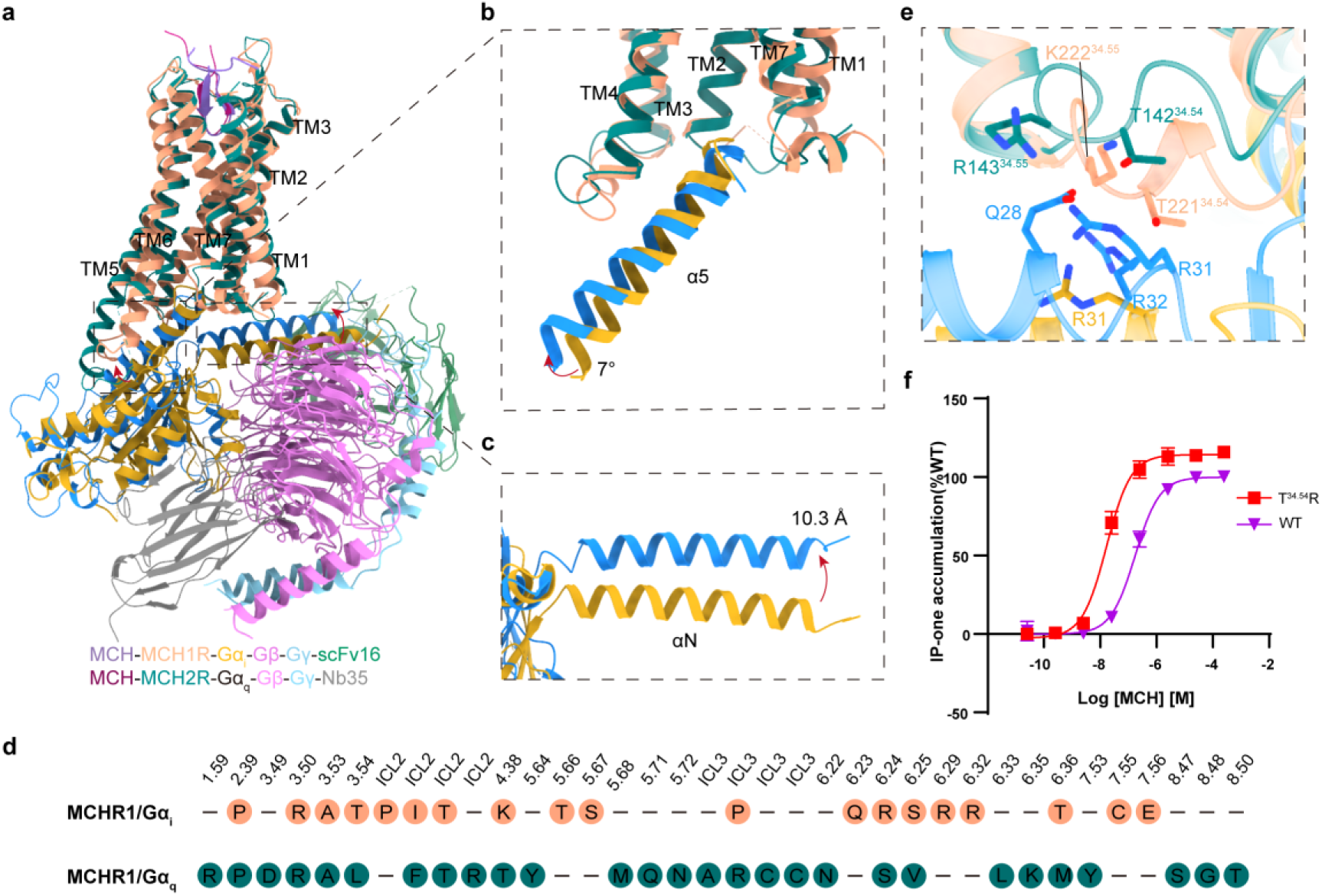
G-Protein coupling of MCHR1 and MCHR2. **a**, Structural alignment between MCHR1-G_i_ and MCHR2-G_q_ complexes. **b**,**c**, Comparative analysis of the Gα conformation in the α5 (**b**) and αN(**c**) domains. **d**, Key residues in MCHR1 and MCHR2 that interact with the downstream Gα_i_ or Gα_q_ proteins. **e**, Detailed interactions between the ICL2 of MCHR1 and MCHR2 with the αN domain of Gα_i_ or Gα_q_. **f**, IP-one assay data for wild-type and T221^34.54^R of MCHR1, with data presented as mean values ± S.E.M. from three independent experiments performed in triplicate.

**Fig. 7.**
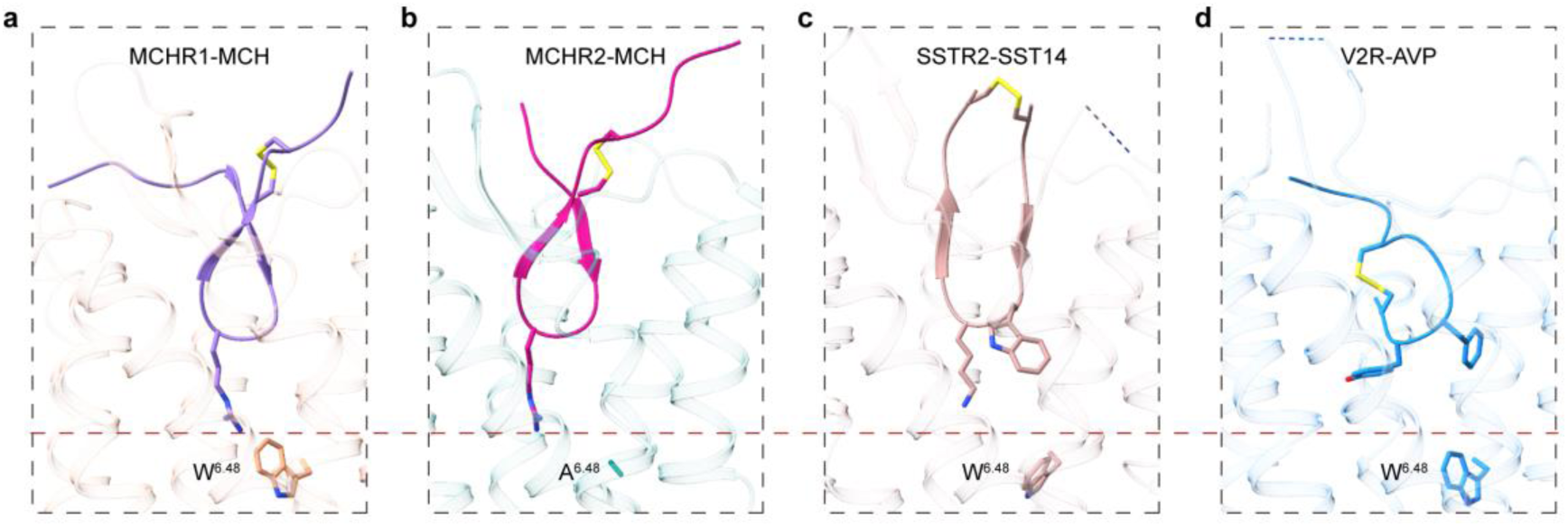
Distinct binding modes of cyclic peptides in receptors. **a**, Binding Conformation of MCH in MCHR1. **b**, Binding Conformation of MCH in MCHR2. **c**, Binding Conformation of SST14 in SSTR2 (PDB: 7Y27). **d**, Binding Conformation of AVP in V2R (PDB: 7DW9).

A salient feature of most Gα_q_-coupled receptors is the incorporation of a basic residue, either arginine or lysine, at the loci 34.54 or 34.55 within ICL2 ^41^. In the context of MCHR1, the threonine residue at position 34.54 (T221) establishes a polar interaction with the αN domain of Gα_q_ (Fig. 6e). A mutation of this threonine to arginine culminated in a tenfold amplification in the coupling efficiency of MCH1R to Gα_q_, thereby accentuating the pivotal role of this residue in G-protein coupling specificity (Fig. 6f). Together, the derived structural insights above offer a comprehensive understanding of the G-protein coupling specificities between MCHR1 and MCHR2.

## Comparison with ligand-activated SSTR2 and V2R

MCH, SST14, and AVP are cyclic polypeptides, each stabilized by disulfide bonds. These peptides adopt analogous conformations, with their cyclic segments deeply embedded within their respective receptor binding pockets (SST14 in SSTR2; AVP in V2R). Distinctively, MCH adopts a twisted γ-shaped ring conformation, diverging from the near-planar cyclic amino acid backbones exhibited by SST14 and AVP (Fig. 7).

SST14, characterized by its expansive loop encompassing 12 amino acids, positions its disulfide bond above the receptor pocket. Central residues, W8 and K9, from its FWKT motif, play pivotal roles in receptor engagement (Fig. 7c). In contrast, AVP, with its compact 6-amino acid ring, situates its disulfide bond intrinsically within the receptor pocket (Fig. 7d). Key residues, Y2 and F3, delve into the pocket’s depths, with their side chains docked horizontally into the pocket, which contrasts with the vertical docking of the sidechain of R11 from MCH into both MCHR1 and MCHR2 pocket (Fig. 7a-b,d). The profound penetration of MCH into the receptor pocket surpasses that of both SST14 and AVP.

## Discussion

In this study, we have elucidated the structural mechanisms of MCHR1-G_i_ and MCHR2-G_q_ upon activation by the endogenous ligand, MCH. Through a combination of high-resolution structural determination, mutagenesis, and functional assays, we have delineated the ligand-binding sites of both MCHR1 and MCHR2. Our findings underscore the significance of the ring structure, formed by Cys6 and Cys17, in facilitating ligand-receptor interactions.

A comparative analysis of the ligand-binding pockets of MCHR1 and MCHR2 revealed distinct molecular interactions. Specifically, S^7.42^ in MCHR2 forms a hydrogen bond with R11, a feature absence in MCHR1 due to the presence of G^7.42^. Additionally, the steric hindrance introduced by W^6.48^ in MCHR1, contrasted with A^6.48^ in MCHR2, results in the displacement of R11 away from F^2.53^ in MCHR1. In MCHR2, H^2.53^ establishes a robust polar interaction with R11. Furthermore, the spatial arrangement of W^2.64^ and Y^7.32^ in MCHR2 positions MCH closer to ECL2, facilitating unique contacts of MCH to MCHR2.

Upon activation, the canonical motifs DRY, NPxxY, and PIF in both MCHR1 and MCHR2 adopt conformations indicative of receptor activation. However, the toggle switch mechanism, typically observed in class A GPCRs, presents a divergence. In MCHR1, W^6.48^ remains relatively static post-activation, while MCHR2 features an alanine at the corresponding position. The binding of MCH disrupts the polar interaction between Q^3.36^ and H/Y^6.52^, leading to conformational shifts in TM6 and TM7.

Our structural insights also shed light on the intracellular interactions between MCHRs and G proteins. Through mutagenesis studies, we identified T^34.54^R in ICL2 of MCHR1 as a critical residue enhancing Gα_q_ response, suggesting that basic residues at this position are conducive for Gα_q_ coupling specificity.

In summary, our findings provide a comprehensive understanding of the molecular mechanisms governing ligand binding, receptor activation, and G protein coupling in MCHRs, offering a foundation for the rational design of therapeutics targeting these receptors.

## Acknowledgements

The cryo-EM data were collected at Advanced Center for Electron Microscopy at Shanghai Institute of Materia Medica, Chinese Academy of Sciences. This work was supported by Natural Science Foundation of Shanghai (23ZR1475200 to L.H.Z); CAS StrateGic Priority Research Program (XDB37030103 to H.E.X.); National Natural Science Foundation of China (32071203 to L.H.Z, 32130022 and 82121005 to H.E.X.); Shanghai Municipal Science and Technology Major Project (2019SHZDZX02 to H.E.X.); Shanghai Municipal Science and Technology Major Project (H.E.X.);the Young Innovator Association of CAS (2018325 and Y2022078 to L.H.Z); the National Key R&D Program of China (2018YFA0507002 to H.E.X., 2019YFA0904200); the Lingang Laboratory (LG-GG-202204-01to H.E.X.); State Key Laboratory of Drug Research (SKLDR-2023-TT-04 to H.E.X.).

## Author contributions

Q.H. and L.H.Z designed the expression constructs, Q.H. purified the complexes, prepared the final samples for cryo-EM data collection toward the structure; Q.H. and L.H.Z prepared the cryo-EM grids, Q.H. performed function assays and prepared figures and drafted manuscript; K.W. and W.H. collected the cryo-EM data, Q.N.Y performed map calculations, Q.N.Y and H.S. built and refined the structure models; C.R.W, Y.M.G and Y.M.Z participated in data analysis; L.H.Z and H.E.X. performed structure and function data analysis, and wrote the manuscript; L.H.Z and H.E.X. conceived, designed, and supervised the overall project.

## Competing interests

The authors declare no competing interests.

## Data availability

Cryo-EM maps have been deposited in the Electron Microscopy Data Bank under accession codes: EMD-XXXXX (MCH-MCHR1-G_i_ complex) and EMD-XXXX (MCH-MCHR2-G_q_ complex). The atomic coordinates have been deposited in the Protein Data Bank under accession codes: XXXX (MCH-MCHR1-G_i_ complex) and 8HAO (MCH-MCHR2-G_q_ complex).

## Code availability

This paper does not report original code.

## Methods

### Constructs cloning

The gene sequences of human MCHR1(residues 70-405) and MCHR2(residues 1-331, with L256^6.40^Y mutation) was subcloned into pFastBac vector (Invitrogen) with an N-terminal haemagglutinin signal peptide (HA) and thermostabilized b562RIL (BRIL), followed by a C-terminal 15-amino acid linker, a LgBiT subunit and two maltose-binding proteins (MBPs) to facilitate receptor expression and stability. The Gα_i_ construct was designed with dominant-negative mutations of S47N, G203, A326, and E245A to decrease the affinity of nucleotide-binding and stable Gαβγ complex. The mini-Gαq construct was designed based on mini-Gα_s_ skeleton and connected with the N-terminus of Gα_i_. Rat Gβ1 was constructed with a N-terminal 16x His-tag for purification and a C-terminal HiBiT to form a NanoBiT with LgBiT. Gα_i_, mini-Gα_q_, Gβ1, Gγ2, scFv16 and Nb35 were all cloned into the pFastBac vector separately. The wild-type and mutants of MCHR1 and MCHR2 were constructed into the pcDNA6.0 vector (Promega) for functional assays.

### Complex expression and purification

MCHRs and G proteins were co-expressed in High Five insect cells (Thermo Fisher) using Bac-to-Bac baculovirus system (Thermo Fisher). The cells were cultured in ESF 921 serum-free medium (Expression Systems) to a density of 3×10^6^ cells /mL. The cells were infected with MCHR1, Gα_i_, Gβ_1_, Gγ_2_, scFv16, and Ric8B at the ratio of 1:2:2:2:2:2 or infected with MCHR2, Gα_q_, Gβ_1_, Gγ_2_, and Ric8A at the ratio of 1:2:2:2:2. After 48 h of infection at 27 ℃, the cells were harvested by centrifugation at 2000 rpm and then stored at -80 ℃ for future use.

The frozen cells were thawed at room temperature (RT) and resuspended in 20 mM HEPES pH 7.4, 100 mM NaCl, 10% (v/v) glycerol, 10 mM MgCl_2_, 10 mM CaCl_2_, 100 μM TCEP, 25 mU/mL apyrase, 1x protease inhibitor cocktail (TargetMol, 1 mL/100 mL suspension), 10 μM MCH and 10 μg/mL Nb35(only for MCHR2). The suspension was incubated at RT for 1h and then solubilized by 0.5% (w/v) lauryl maltose neopentyl glycol (LMNG, Anatrace) and 0.1% (w/v) cholesteryl hemisuccinate TRIS salt (CHS, Anatrace) at 4°C for 3 h. The supernatant was separated by centrifugation at 30000 rpm for 35 min and then incubated with re-equilibrated dextrin beads (Smart-Lifesciences) at 4 ℃ for 3 h. The resin was collected by centrifugation at 500× g for 5 min, loaded onto a gravity-flow column and washed with 20 column volumes of wash buffer containing 20 mM HEPES pH 7.4, 100 mM NaCl, 10% (v/v) glycerol, 5 mM MgCl_2_, 5 mM CaCl_2_, 25 μM TCEP, 10 μM MCH, 0.01% (w/v) LMNG, 0.01% (w/v) glyco-diosgenin (GDN, Anatrace) and 0.004% (w/v) CHS. The protein was eluted with wash buffer adding 10 mM maltose, concentrated using a 100-kD Amicon Ultra Centrifugal Filter (Millipore) and loaded onto Superose 6 Increase 10/300 GL column (GE Healthcare) pre-equilibrated with buffer including 20 mM HEPES pH 7.4, 100 mM NaCl, 2 mM MgCl_2_, 2 mM CaCl_2_, 100 μM TCEP, 0.0005 % (w/v) digitonin (Biosynth), 0.00075% (w/v) LMNG, 0.00025% (w/v) GDN, and 0.0002% (w/v) CHS. The receptor and G protein complexes were collected and concentrated for electron microscopy experiments.

### Expression and purification of Nb35

Nanobody-35 (Nb35) was expressed in *E. coli* BL21 cells which were cultured in TB medium with 100 μg/mL ampicillin at 37 ℃ for about 3h to OD600 of 1.0 and then induced with 1mM IPTG at 25℃ for 16h. Each liter of harvested cells was resuspended with 15mL TES (0.2 M Tris pH 8.0, 0.5 mM EDTA, 0.5 M sucrose), stirred at 4 ℃ for 1h and another 45 min with 30 mL TES/4. The supernatant was collected by centrifugation at 30000 rpm for 30 min and incubated with pre-equilibrated Nickel resin at 4 ℃ for 1h. After washing with 20 column volumes of wash buffer containing 20 mM HEPES pH 7.4, 100 mM NaCl, 10% (v/v) glycerol, Nb35 was eluted with wash buffer adding 300 mM imidazole. The protein was further purified by HiLoad 16/600 Superdex 75 column with 20 mM HEPES pH 7.4, 100 mM NaCl. The targeted fractions were collected and concentrated to 5mg/mL, and then flash-frozen in liquid nitrogen before storing at -80 °C.

### Cryo-EM grid preparation and data collection

For cryo-EM grid preparation of the MCH-MCHR1-G_i_ complex, 3μL of the purified complex at a concentration of 5.72 mg/mL was applied to glow-discharged holey carbon grids (Quantifoil R1.2/1.3, Au/C 300 mesh) that were subsequently vitrified by plunging into liquid ethane using a Vitrobot Mark IV (ThermoFisher Scientific) at 4 ℃. A Titan Krios G4 equipped with a Gatan K3 direct electron detector with super-resolution mode and EPU were used to acquire cryo-EM movies at Advanced Center for Electron Microscopy at Shanghai Institute of Materia Medica, Chinese Academy of Sciences. A total of 14,952 Movies were recorded with pixel size of 0.824 Å at a dose of 50 electron per Å^2^ for 32 frames. The defocus range of this dataset was -0.8 μm to -1.8 μm.

For the MCH-MCHR2-G_q_ complex, 20.47 mg/mL of purified protein was used for cryo-EM grid preparation. Cryo-EM movies were collected by a Titan Krios G4 CEFG at 300KV accelerating voltage equipped with Falcon4 direct electron detector and Selectris X (ThermoFisher Scientific). A total of 8,789 EER movies were recorded with pixel size of 0.73 Å at a dose of 50 electron per Å^2^. The defocus range of this dataset was -0.8 μm to -1.8 μm.

### Cryo-EM data processing three-dimensional reconstruction

For the MCH-MCHR1-G_i_ complex, all dose-fractionated image stacks were subjected to beam-induced motion correction by Relion4.0^42^. The defocus parameters were estimated by CTFFIND4.1^43^. Blob picking and Template picking yielded 32,120,445 particles, which were processed by reference-free 2D classification using Cryosparc ^44^. With initial model from Cryosparc, after several rounds of 3D classification using Relion, 311,033 particles were used to further CTF Refinement and polishing, yielding a reconstruction with global resolution of 3.01Å at a Fourier shell correlation (FSC) of 0.143, and subsequently post-processed by DeepEMhancer^45^.

For the MCH-MCHR2-G_q_ complex, EER movies were aligned with Relion4.0^42^. Initial contrast transfer function(CTF) fitting was performed with CTFFIND4.1^43^ from Cryosparc^44^. Blob picking and Template picking yielded 14,206,080 particles, which were processed by reference-free 2D classification using Cryosparc, 2D Classification were processed using Cryosparc, producing 1,498,863 particles for further processing. With initial model, two rounds of 3D classifications were carried out with Relion4.0 in which 879,702 particles were subjected to 3D auto-refinement, CTF refinement and polishing. A map with an indicated global resolution of 2.4 Å at a Fourier shell correlation (FSC) of 0.143 was generated from the final 3D refinement, and subsequently post-processed by DeepEMhancer^45^.

### Model building and refinement

All PDB coordinates using alphafold2^46^ were served as a starting model for building the atomic model. All model were fitted into cryo-EM density map using chimera ^47^ followed by a manual adjustment in Coot^48^. The model was refined by Phenix ^49^.

### GloSensor cAMP assay

GloSensor cAMP assay was used to detecte the downstream Gα_i_ signal of MCHR1 using GloSensor cAMP Reagent (Promega). HEK293 cells were cultured in DMEM/ high Glucose supplemented with 10% (v/v) fetal bovine serum and 1% (v/v) Penicillin-Streptomycin at 37 °C in 5% CO_2_. Cells was grown at 12-well plates for 18h and transfected with MCHR1 constructs and the cAMP biosensor GloSensor-22F (Promega) at a ratio of 3:2. After transfection for 24h, cells were harvested, resuspended with CO_2_-independent media containing 2% GloSensor cAMP Reagent, and then distributed to 384-well plates at a density of 3×10^5^ cells/mL (20 μL/well). The cells were incubated at 37 °C for 1.5h. Ligand of different concentrations was mixed with forskolin at the final concentration of 1 μM (Sigma) and the mixture was added to each well (10 μL /well). The sample mixtures were immediately measured with EnVision multi-plate reader (PerkinElmer).

### Inositol phosphate accumulation assay

IP1 accumulation assay was applied for detection of the downstream Gα_q_ signal of MCHR2 using the IP-One HTRF kit (Cisbio). AD293 cells were cultured in DMEM/ high Glucose supplemented with 10% (v/v) fetal bovine serum and 1% (v/v) Penicillin-Streptomycin at 37 °C in 5% CO_2_. After transfection for 24h, cells were harvested, resuspended with 1× stimulation buffer to a density of 8 × 10^5^ cells/mL, and then seeded to 384-well plates for 7 μL/well. After dispensing 7 μL different concentrations of ligand diluted with 1× stimulation buffer, the mixture was incubated at 37 °C for 1h. IP1-d2 and anti-IP1 cryptate was dissolved in 1× lysis buffer and sequentially added to the plates for 3 μL/well. Before measurement, the samples were incubated at RT for 30 min and measured with EnVision multi-plate reader (PerkinElmer).

### Quantification and statistical analysis

All functional study data were analyzed using Prism 8 (GraphPad) and displayed as means ± S.E.M. from at least three independent experiments. Concentration-response curves were evaluated with a three-parameter logistic equation. The significance was determined with one-way ANOVA with Tukey’s test, and *P* < 0.05 vs. wild type (WT) was considered statistically significant.

## Notes

### Competing Interest Statement

The authors have declared no competing interest.

## References

1 Kawauchi, H., Kawazoe, I., Tsubokawa, M., Kishida, M. & Baker, B. I. Characterization of melanin-concentrating hormone in chum salmon pituitaries. Nature 305, 321–323, doi:10.1038/305321a0 (1983).

2 Skofitsch, G., Jacobowitz, D. M. & Zamir, N. Immunohistochemical localization of a melanin concentrating hormone-like peptide in the rat brain. Brain Res Bull 15, 635–649, doi:10.1016/0361-9230(85)90213-8 (1985).

3 Vaughan, J. M., Fischer, W. H., Hoeger, C., Rivier, J. & Vale, W. Characterization of melanin-concentrating hormone from rat hypothalamus. Endocrinology 125, 1660–1665, doi:10.1210/endo-125-3-1660 (1989).

4 Bittencourt, J. C. et al. The melanin-concentrating hormone system of the rat brain: an immuno- and hybridization histochemical characterization. J Comp Neurol 319, 218–245, doi:10.1002/cne.903190204 (1992).

5 Saito, Y., Nothacker, H. P. & Civelli, O. Melanin-concentrating hormone receptor: an orphan receptor fits the key. Trends Endocrinol Metab 11, 299–303, doi:10.1016/s1043-2760(00)00290-3 (2000).

6 Nahon, J. L. The melanocortins and melanin-concentrating hormone in the central regulation of feeding behavior and energy homeostasis. C R Biol 329, 623–638; discussion 653-625, doi:10.1016/j.crvi.2006.03.021 (2006).

7 Al-Massadi, O. et al. Multifaceted actions of melanin-concentrating hormone on mammalian energy homeostasis. Nat Rev Endocrinol 17, 745–755, doi:10.1038/s41574-021-00559-1 (2021).

8 Jiang, H. & Bruning, J. C. Melanin-Concentrating Hormone-Dependent Control of Feeding: When Volume Matters. Cell Metab 28, 7–8, doi:10.1016/j.cmet.2018.06.018 (2018).

9 Bandaru, S. S., Khanday, M. A., Ibrahim, N., Naganuma, F. & Vetrivelan, R. Sleep-Wake Control by Melanin-Concentrating Hormone (MCH) Neurons: a Review of Recent Findings. Curr Neurol Neurosci Rep 20, 55, doi:10.1007/s11910-020-01075-x (2020).

10 Torterolo, P. et al. Melanin-Concentrating Hormone (MCH): Role in REM Sleep and Depression. Front Neurosci 9, 475, doi:10.3389/fnins.2015.00475 (2015).

11 Adamantidis, A. et al. Disrupting the melanin-concentrating hormone receptor 1 in mice leads to cognitive deficits and alterations of NMDA receptor function. European Journal of Neuroscience 21, 2837–2844, doi:10.1111/j.1460-9568.2005.04100.x (2005).

12 Tyhon, A. et al. Mice lacking the melanin-concentrating hormone receptor-1 exhibit an atypical psychomotor susceptibility to cocaine and no conditioned cocaine response. Behav Brain Res 173, 94–103, doi:10.1016/j.bbr.2006.06.007 (2006).

13 Schneider, J. E. Energy balance and reproduction. Physiol Behav 81, 289–317, doi:10.1016/j.physbeh.2004.02.007 (2004).

14 Naufahu, J., Cunliffe, A. D. & Murray, J. F. The roles of melanin-concentrating hormone in energy balance and reproductive function: Are they connected? Reproduction 146, R141–150, doi:10.1530/REP-12-0385 (2013).

15 Prida, E. et al. Crosstalk between Melanin Concentrating Hormone and Endocrine Factors: Implications for Obesity. International Journal of Molecular Sciences 23, 2436 (2022).

16 Hausen, A. C. et al. Insulin-Dependent Activation of MCH Neurons Impairs Locomotor Activity and Insulin Sensitivity in Obesity. Cell Rep 17, 2512–2521, doi:10.1016/j.celrep.2016.11.030 (2016).

17 Roy, M., David, N., Cueva, M. & Giorgetti, M. A study of the involvement of melanin-concentrating hormone receptor 1 (MCHR1) in murine models of depression. Biol Psychiatry 61, 174–180, doi:10.1016/j.biopsych.2006.03.076 (2007).

18 Roy, M. et al. Genetic inactivation of melanin-concentrating hormone receptor subtype 1 (MCHR1) in mice exerts anxiolytic-like behavioral effects. Neuropsychopharmacology 31, 112–120, doi:10.1038/sj.npp.1300805 (2006).

19 Potter, L. E. & Burgess, C. R. The melanin-concentrating hormone system as a target for the treatment of sleep disorders. Front Neurosci 16, 952275, doi:10.3389/fnins.2022.952275 (2022).

20 Jeon, J. Y. et al. MCH-/-mice are resistant to aging-associated increases in body weight and insulin resistance. Diabetes 55, 428–434, doi:10.2337/diabetes.55.02.06.db05-0203 (2006).

21 Willie, J. T., Sinton, C. M., Maratos-Flier, E. & Yanagisawa, M. Abnormal response of melanin-concentrating hormone deficient mice to fasting: hyperactivity and rapid eye movement sleep suppression. Neuroscience 156, 819–829, doi:10.1016/j.neuroscience.2008.08.048 (2008).

22 Bell, C. G. et al. Association of melanin-concentrating hormone receptor 1 5’ polymorphism with early-onset extreme obesity. Diabetes 54, 3049–3055, doi:10.2337/diabetes.54.10.3049 (2005).

23 Ahnaou, A., Dautzenberg, F. M., Huysmans, H., Steckler, T. & Drinkenburg, W. H. Contribution of melanin-concentrating hormone (MCH1) receptor to thermoregulation and sleep stabilization: evidence from MCH1 (-/-) mice. Behav Brain Res 218, 42–50, doi:10.1016/j.bbr.2010.11.019 (2011).

24 Tan, C. P. et al. Melanin-concentrating hormone receptor subtypes 1 and 2: species-specific gene expression. Genomics 79, 785–792, doi:10.1006/geno.2002.6771 (2002).

25 Saito, Y. et al. Molecular characterization of the melanin-concentrating-hormone receptor. Nature 400, 265–269, doi:10.1038/22321 (1999).

26 Hawes, B. E. et al. The Melanin-Concentrating Hormone Receptor Couples to Multiple G Proteins to Activate Diverse Intracellular Signaling Pathways. Endocrinology 141, 4524–4532, doi:10.1210/endo.141.12.7833 (2000).

27 Pissios, P., Trombly, D. J., Tzameli, I. & Maratos-Flier, E. Melanin-concentrating hormone receptor 1 activates extracellular signal-regulated kinase and synergizes with G(s)-coupled pathways. Endocrinology 144, 3514–3523, doi:10.1210/en.2002-0004 (2003).

28 Sailer, A. W. et al. Identification and characterization of a second melanin-concentrating hormone receptor, MCH-2R. Proceedings of the National Academy of Sciences 98, 7564–7569, doi:doi:10.1073/pnas.121170598 (2001).

29 Baker, B. I. Melanin-concentrating hormone: a general vertebrate neuropeptide. Int Rev Cytol 126, 1–47, doi:10.1016/s0074-7696(08)60681-6 (1991).

30 Chung, S., Parks, G. S., Lee, C. & Civelli, O. Recent updates on the melanin-concentrating hormone (MCH) and its receptor system: lessons from MCH1R antagonists. J Mol Neurosci 43, 115–121, doi:10.1007/s12031-010-9411-4 (2011).

31 Cheon, H. G. Antiobesity effects of melanin-concentrating hormone receptor 1 (MCH-R1) antagonists. Handb Exp Pharmacol, 383–403, doi:10.1007/978-3-642-24716-3_18 (2012).

32 Borowsky, B. et al. Antidepressant, anxiolytic and anorectic effects of a melanin-concentrating hormone-1 receptor antagonist. Nat Med 8, 825–830, doi:10.1038/nm741 (2002).

33 Johansson, A. & Lofberg, C. Novel MCH1 receptor antagonists: a patent review. Expert Opin Ther Pat 25, 193–207, doi:10.1517/13543776.2014.993382 (2015).

34 Chun, E. et al. Fusion partner toolchest for the stabilization and crystallization of G protein-coupled receptors. Structure 20, 967–976, doi:10.1016/j.str.2012.04.010 (2012).

35 Zhao, L. H. et al. Structure insights into selective coupling of G protein subtypes by a class B G protein-coupled receptor. Nat Commun 13, 6670, doi:10.1038/s41467-022-33851-3 (2022).

36 Zhao, L. H. et al. Conserved class B GPCR activation by a biased intracellular agonist. Nature, doi:10.1038/s41586-023-06467-w (2023).

37 Zhao, L. H. et al. Molecular recognition of two endogenous hormones by the human parathyroid hormone receptor-1. Acta Pharmacol Sin, doi:10.1038/s41401-022-01032-z (2022).

38 Zhao, L. H. et al. Structure and dynamics of the active human parathyroid hormone receptor-1. Science 364, 148–153, doi:10.1126/science.aav7942 (2019).

39 Liang, Y. L. et al. Dominant Negative G Proteins Enhance Formation and Purification of Agonist-GPCR-G Protein Complexes for Structure Determination. ACS Pharmacol Transl Sci 1, 12–20, doi:10.1021/acsptsci.8b00017 (2018).

40 An, S. et al. Identification and characterization of a melanin-concentrating hormone receptor. Proc Natl Acad Sci U S A 98, 7576–7581, doi:10.1073/pnas.131200698 (2001).

41 Duan, J. et al. Molecular basis for allosteric agonism and G protein subtype selectivity of galanin receptors. Nat Commun 13, 1364, doi:10.1038/s41467-022-29072-3 (2022).

42. Zivanov, J., Nakane, T. & Scheres, S. H. W. Estimation of high-order aberrations and anisotropic magnification from cryo-EM data sets in RELION-3.1. IUCrJ 7, 253–267, doi:10.1107/S2052252520000081 (2020).

43 Rohou, A. & Grigorieff, N. CTFFIND4: Fast and accurate defocus estimation from electron micrographs. J Struct Biol 192, 216–221, doi:10.1016/j.jsb.2015.08.008 (2015).

44 Punjani, A., Rubinstein, J. L., Fleet, D. J. & Brubaker, M. A. cryoSPARC: algorithms for rapid unsupervised cryo-EM structure determination. Nat Methods 14, 290–296, doi:10.1038/nmeth.4169 (2017).

45 Sanchez-Garcia, R. et al. DeepEMhancer: a deep learning solution for cryo-EM volume post-processing. Commun Biol 4, 874, doi:10.1038/s42003-021-02399-1 (2021).

46 Senior, A. W. et al. Improved protein structure prediction using potentials from deep learning. Nature 577, 706–710, doi:10.1038/s41586-019-1923-7 (2020).

47 Pettersen, E. F. et al. UCSF Chimera--a visualization system for exploratory research and analysis. J Comput Chem 25, 1605–1612, doi:10.1002/jcc.20084 (2004).

48 Emsley, P. & Cowtan, K. Coot: model-building tools for molecular graphics. Acta Crystallogr D Biol Crystallogr 60, 2126–2132, doi:10.1107/S0907444904019158 (2004).

49 Adams, P. D. et al. PHENIX: a comprehensive Python-based system for macromolecular structure solution. Acta Crystallogr D Biol Crystallogr 66, 213–221, doi:10.1107/S0907444909052925 (2010).

