## Extended Data Fig. 1,Extended Data Fig. 2Extended Data Fig. 3,Extended Data Fig. 4.Extended Data Fig. 5, Table 1, Table 2, Table 3 for "Mechanisms of ligand recognition and activation of melanin-concentrating hormone receptors": MCHRs-Supplemental Text and Figures.pdf

**Table of contents**

**Extended Data Fig. 1 | MCH-MCHR1-$G_i$ complexes purification and cryo-EM data processing.**

**Extended Data Fig. 2 | MCH-MCHR2-$G_q$ complexes purification and cryo-EM data processing.**

**Extended Data Fig. 3 | Local density maps of MCH-MCHR1-$G_i$ and MCH-MCHR2-$G_q$ complexes.**

**Extended Data Fig. 4 | Sequence alignment of MCHR1, MCHR2, SSTR2 and V2R.**

**Extended Data Fig. 5 | Mutagenesis data of key residues in the ligand binding pockets of MCHR1 and MCHR2.**

**Table 1 | Interaction of MCH with key residues in the ligand-binding pockets of MCHR1.**

**Table 2 | Interaction of MCH with key residues in the ligand-binding pockets of MCHR2.**

**Table 3 | Comparison of crucial residues within the ligand-binding pockets between MCHR1 and MCHR2.**

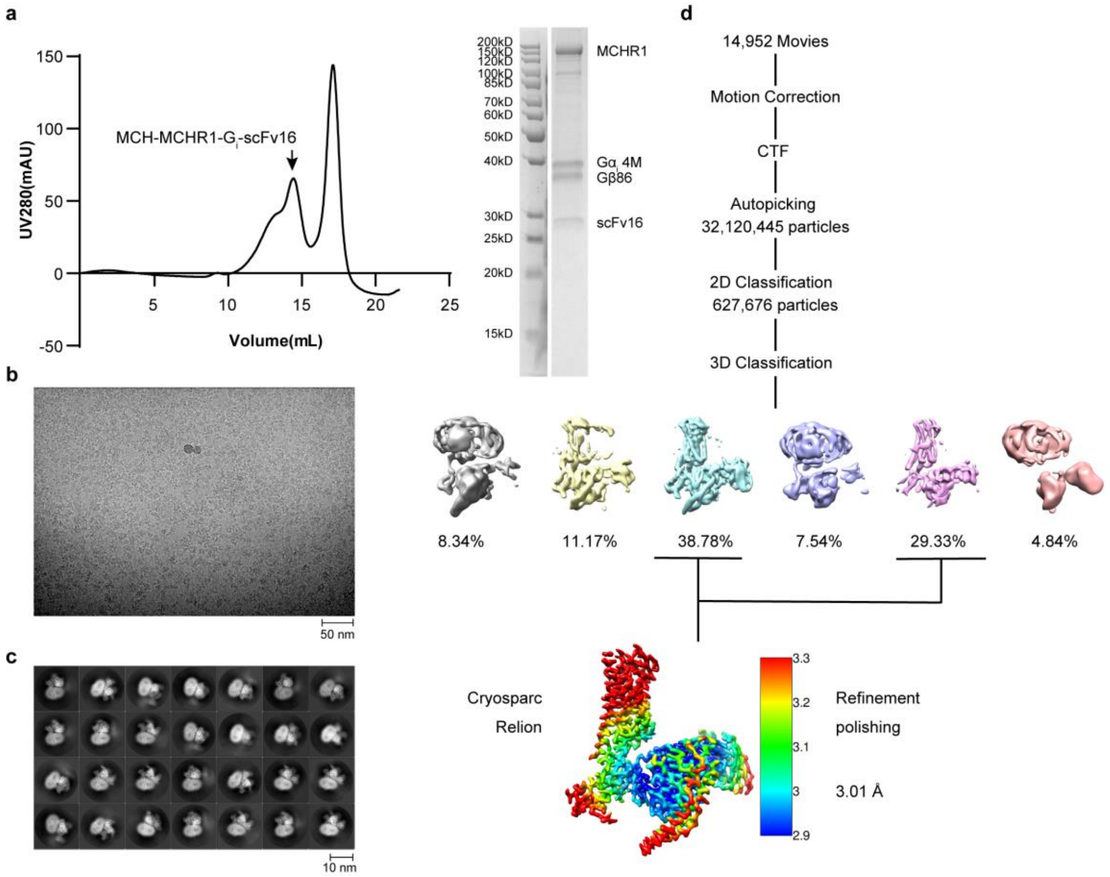


**Extended Data Fig. 1 | MCH-MCHR1-$G_i$ complexes purification and cryo-EM data processing. Related to Figure 1. a**, Profiles of size-exclusion chromatography elution (left) and SDS-PAGE analysis (right) of MCH-MCHR1-$G_i$ complexes. Black arrow refers to complex monomer. **b**,**c**, Representative cryo-EM images (**b**) and 2D classification (**c**) of the MCH-MCHR1-$G_i$ complexes. **d**, Workflow of cryo-EM single particle analysis of the MCH-MCHR1-$G_i$ complexes.

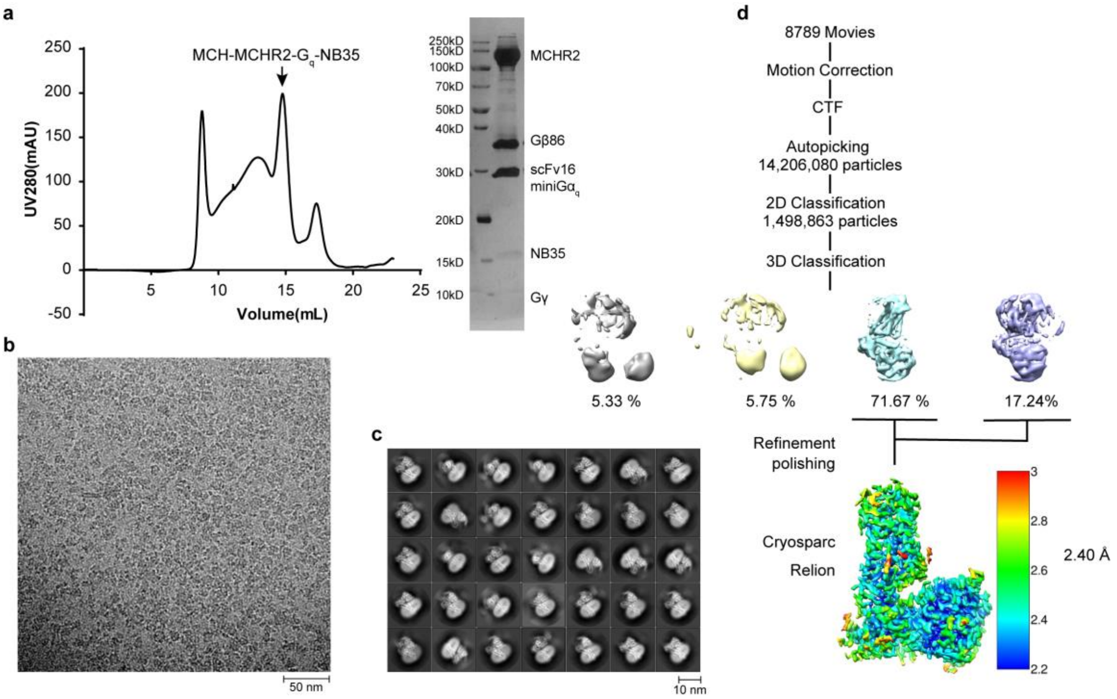


**Extended Data Fig. 2 | MCH-MCHR2-G$_q$ complexes purification and cryo-EM data processing. Related to Figure 1. a**, Profiles of size-exclusion chromatography elution (left) and SDS-PAGE analysis (right) of MCH-MCHR2-G$_q$ complexes. Black arrow refers to complex monomer. **b**,**c**, Representative cryo-EM image (**b**) and 2D classification (**c**) of the MCH-MCHR2-G$_q$ complexes. **d**, Workflow of cryo-EM single particle analysis of the MCH-MCHR2-G$_q$ complexes.

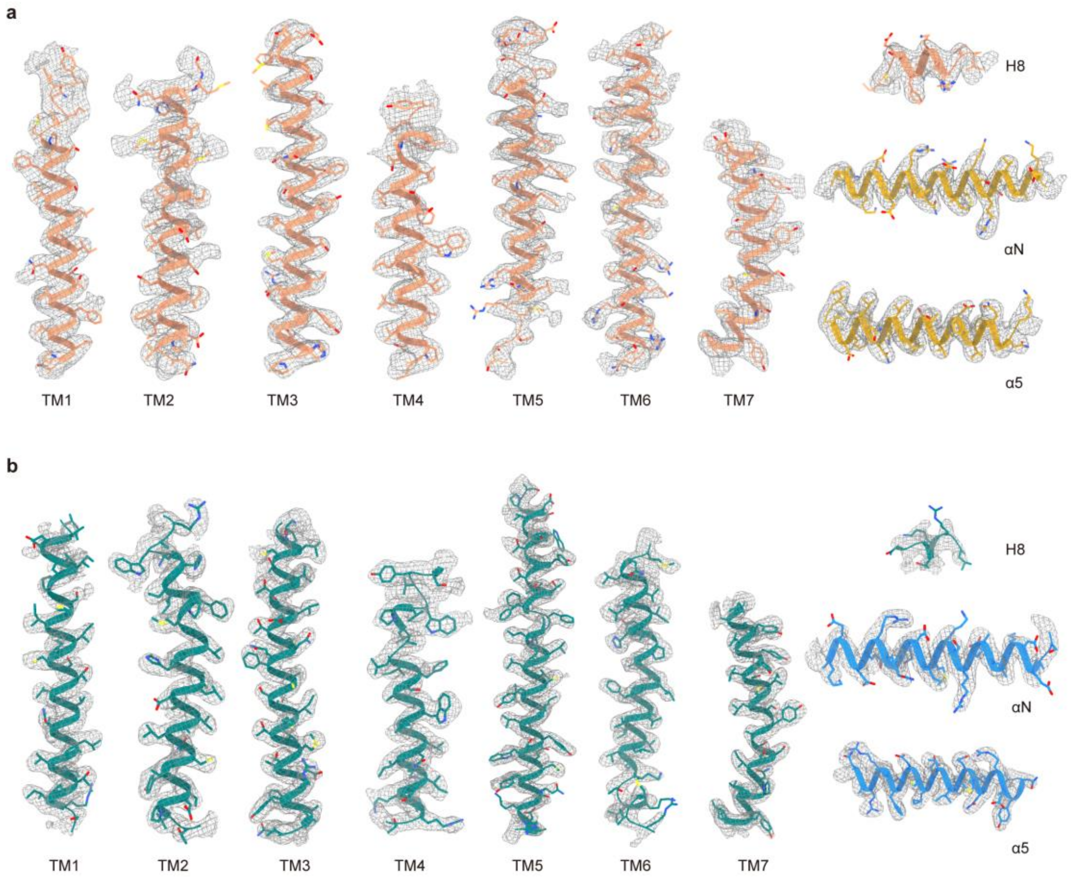


**Extended Data Fig. 3 | Local density maps of MCH-MCHR1-$G_i$ and MCH-MCHR2-$G_q$ complexes. Related to Figure 1. a**, Depicts the representative resolution map of TM1–TM7 and helix 8 of MCHR1, as well as the αN helix and α5 helix of $G\alpha_i$ within the MCH-MCHR1-$G_i$ complexes. **b**, Depicts the representative resolution map of TM1–TM7 and helix 8 of MCHR2, along with the αN helix and α5 helix of $G\alpha_q$ within the MCH-MCHR2-$G_q$ complexes.

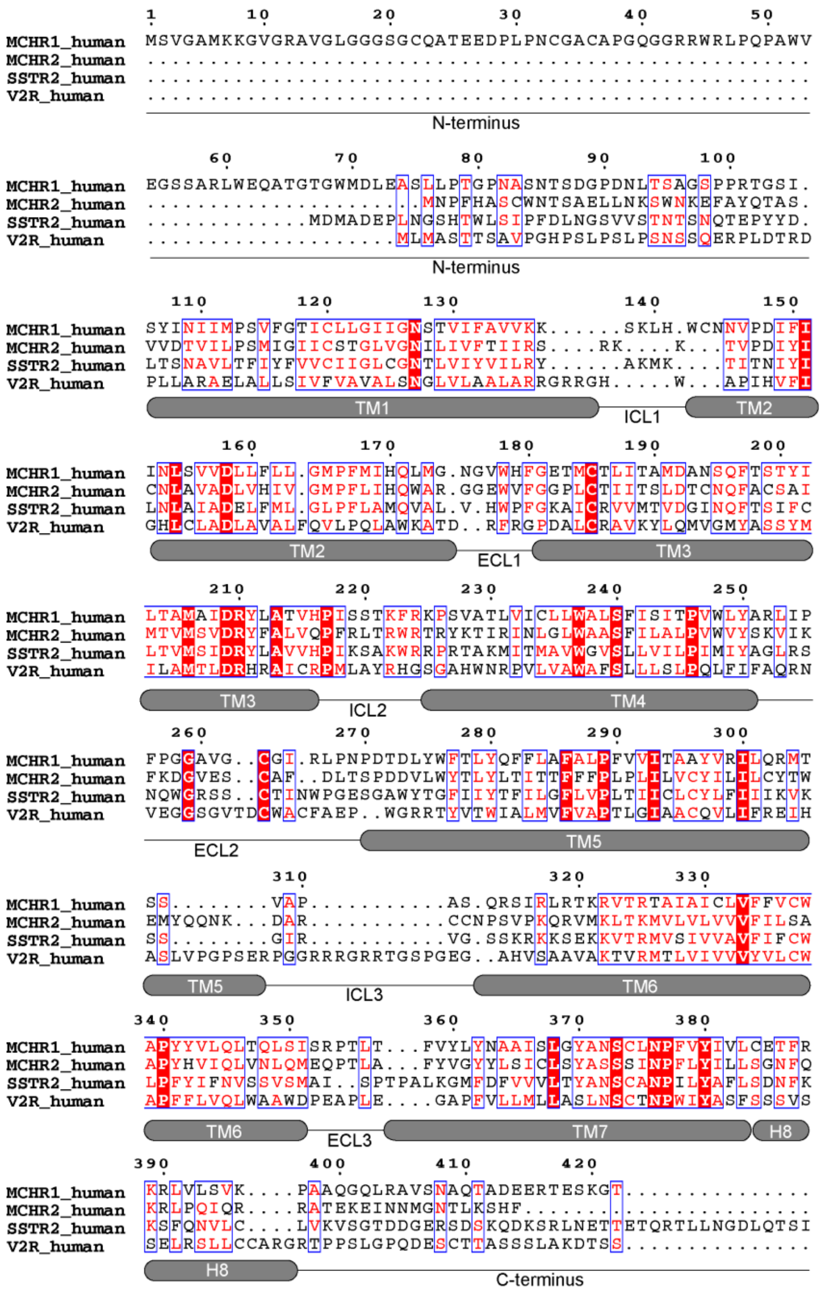


**Extended Data Fig. 4 | Sequence alignment of MCHR1, MCHR2, SSTR2 and V2R. Related to Figure 7.** The sequence alignment of MCHR1, MCHR2, SSTR2 and V2R was created by GPCRdb and the graphic was drawn on the ESPript 3.0 website. TM1-7 and helix8 are shown by columns under the sequences. Colors represent the similarity of residue: red background, identical; red text, strongly similar.

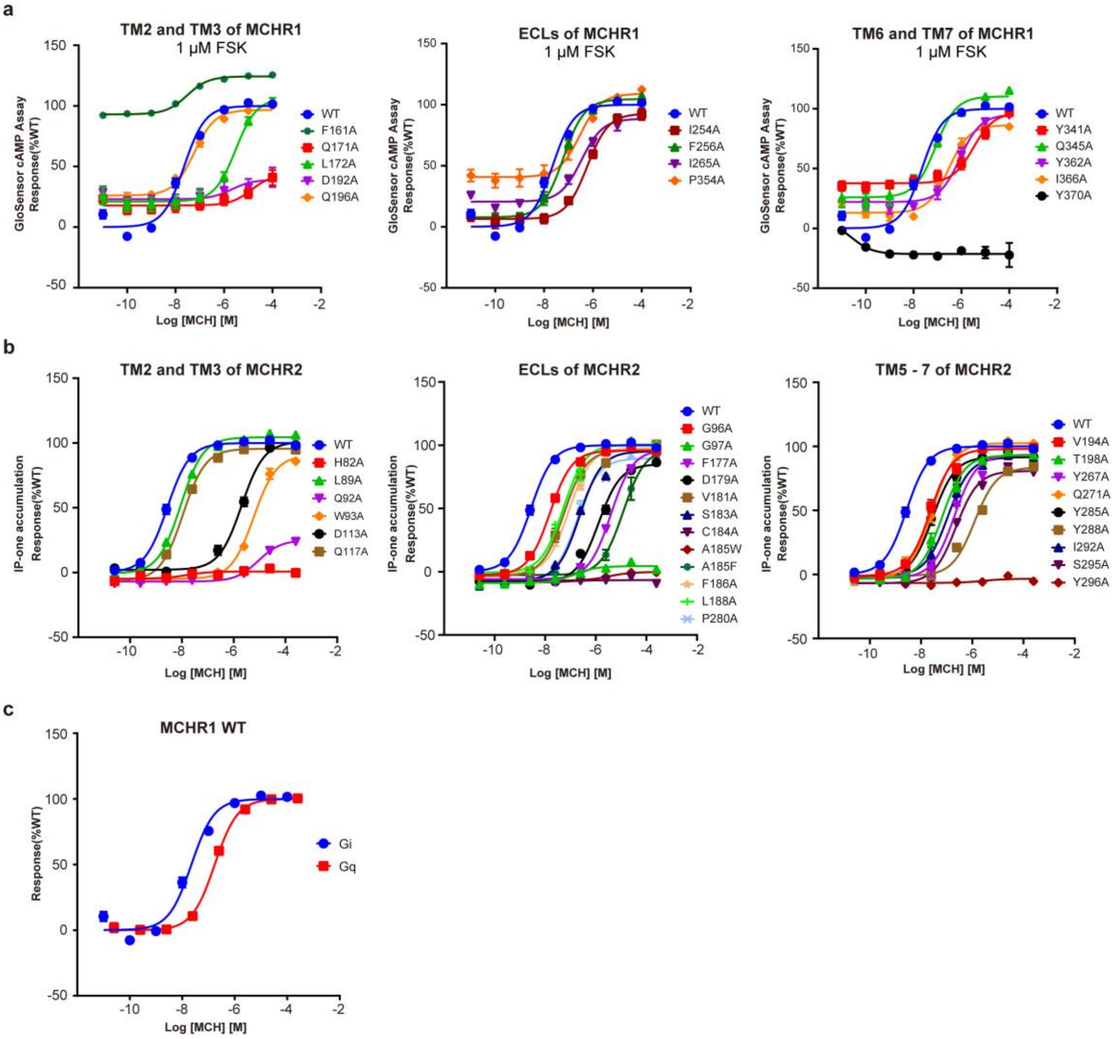


**Extended Data Fig. 5 | Mutagenesis data of key residues in the ligand binding pockets of MCHR1 and MCHR2. Related to Figure 2 and 3. a**, Effects of mutations in the binding pocket of MCHR1 by Glo-sensor cAMP assay. **b**, Effects of mutations in the binding pocket of MCHR2 by IP-one accumulation assay. **c**, Comparison of the efficacy of MCH-activated MCHR1 in coupling with $G_i$ and $G_q$. The response data was normalized by WT receptor within each individual experiment. Data from three independent experiments, each of which was performed in triplicate, are presented as mean ± SEM.

**Table 1 | Interaction of MCH with key residues in the ligand-binding pockets of MCHR1.**

| MCH | MCHR1 | Interactions |
|---|---|---|
| Leu5 | $P354^{ECL3}$ | Hydrophobic interaction |
| Arg6 | | |
| Met8 | $L172^{2.64}$ | Hydrophobic interaction |
| | $Y362^{7.53}$ | |
| Leu9 | $Y362^{7.53}$ | Hydrophobic interaction |
| Gly10 | $Y341^{6.51}$ | Hydrophobic interaction |
| | $Y362^{7.53}$ | |
| Arg11 | $F161^{2.53}$ | Hydrophobic interaction |
| | $D192^{3.32}$ | Electrostatic interaction |
| | $Q196^{3.36}$ | Side chain-side chain hydrogen bond |
| | $I366^{7.39}$ | Hydrophobic interaction |
| | $Y370^{7.43}$ | |
| Val12 | $Q171^{2.63}$ | Backbone-side chain hydrogen bond |
| | $I265^{ECL2}$ | Hydrophobic interaction |
| Tyr13 | $F256^{ECL2}$ | Hydrophobic interaction |
| Pro15 | $I254^{ECL2}$ | Hydrophobic interaction |

**Table 2 | Interaction of MCH with key residues in the ligand-binding pockets of MCHR2.**

| MCH | MCHR2 | Interactions |
|---|---|---|
| Leu5 | $F177^{ECL2}$ | Hydrophobic interaction |
| Arg6 | $D179^{ECL2}$ | Electrostatic interaction |
| | $V181^{ECL2}$ | Hydrophobic interaction |
| | $G96^{ECL1}$ | |
| Met8 | $W93^{2.64}$ | Hydrophobic interaction |
| | $P280^{ECL3}$ | |
| | $Y285^{7.32}$ | |
| Leu9 | $F186^{ECL2}$ | Hydrophobic interaction |
| | $L188^{ECL2}$ | |
| | $V194^{ECL2}$ | |
| | $T198^{ECL2}$ | |
| | $Y288^{7.35}$ | |
| Gly10 | $Q271^{6.55}$ | Backbone-side chain hydrogen bond |
| | $Y267^{6.51}$ | Hydrophobic interaction |
| Arg11 | $H82^{2.53}$ | Hydrophobic interaction |
| | $W93^{2.64}$ | Backbone-side chain hydrogen bond |
| | $D113^{3.32}$ | Electrostatic interaction |
| | $Q117^{3.36}$ | |
| | $Y288^{7.35}$ | Hydrophobic interaction |
| | $I292^{7.39}$ | |
| | $S295^{7.42}$ | Side chain-side chain hydrogen bond |
| | $Y296^{7.43}$ | Hydrophobic interaction |
| Vla12 | $L89^{2.60}$ | Hydrophobic interaction |
| | $C184^{ECL2}$ | |
| | $F186^{ECL2}$ | |
| Tyr13 | $Q92^{2.63}$ | Hydrophobic interaction |
| | $W93^{2.64}$ | |
| | $G97^{ECL1}$ | |
| | $S183^{ECL2}$ | |
| | $C184^{ECL2}$ | |
| | $A185^{ECL2}$ | |
| | $F186^{ECL2}$ | Backbone-backbone hydrogen bond |
| Pro15 | $F186^{ECL2}$ | Hydrophobic interaction |
| | $L188^{ECL2}$ | |

**Table 3 | Comparison of crucial residues within the ligand-binding pockets between MCHR1 and MCHR2.**

| MCH | MCHR1 | MCHR2 |
|---|---|---|
| Leu5 | P354$^{ECL3}$ | F177$^{ECL2}$ |
| Arg6 | P354$^{ECL3}$ | D179$^{ECL2}$ |
| | | V181$^{ECL2}$ |
| | | G96$^{ECL1}$ |
| Met8 | L172$^{2.64}$ | W93$^{2.64}$ |
| | | P280$^{ECL3}$ |
| | Y362$^{7.35}$ | Y285$^{7.32}$ |
| Leu9 | | F186$^{ECL2}$ |
| | | L188$^{ECL2}$ |
| | | V194$^{ECL2}$ |
| | | T198$^{ECL2}$ |
| | Y362$^{7.35}$ | Y288$^{7.35}$ |
| Gly10 | Y341$^{6.51}$ | Y267$^{6.51}$ |
| | Y362$^{7.53}$ | Q271$^{6.55}$ |
| Arg11 | F161$^{2.53}$ | H82$^{2.53}$ |
| | | W93$^{2.64}$ |
| | D192$^{3.32}$ | D113$^{3.32}$ |
| | Q196$^{3.36}$ | Q117$^{3.36}$ |
| | | Y288$^{7.35}$ |
| | I366$^{7.39}$ | I292$^{7.39}$ |
| | | S295$^{7.42}$ |
| | Y370$^{7.43}$ | Y296$^{7.43}$ |
| Val12 | | L89$^{2.60}$ |
| | | C184$^{ECL2}$ |
| | I265$^{ECL2}$ | F186$^{ECL2}$ |
| Tyr13 | F256$^{ECL2}$ | Q92$^{2.63}$ |
| | | W93$^{2.64}$ |
| | | G97$^{ECL1}$ |
| | | S183$^{ECL2}$ |
| | | C184$^{ECL2}$ |
| | | A185$^{ECL2}$ |
| | | F186$^{ECL2}$ |
| Pro15 | I254$^{ECL2}$ | F186$^{ECL2}$ |
| | | L188$^{ECL2}$ |
